# Functional redundancy of variant and canonical histone H3 lysine 9 modification in *Drosophila*

**DOI:** 10.1101/183251

**Authors:** Taylor J.R. Penke, Daniel J. McKay, Brian D. Strahl, A. Gregory Matera, Robert J. Duronio

**Author notes:** To whom correspondence should be addressed, Robert J. Duronio Integrative Program for Biological and Genome Sciences, CB#7100, University of North Carolina Chapel Hill, NC 27599, Voice.

## Abstract

Histone post-translational modifications (PTMs) and differential incorporation of variant and canonical histones into chromatin are central modes of epigenetic regulation. Despite similar protein sequences, histone variants are enriched for different suites of PTMs compared to their canonical counterparts. For example, variant histone H3.3 occurs primarily in transcribed regions and is enriched for “active” histone PTMs like Lys9 acetylation (H3.3K9ac), whereas the canonical histone H3 is enriched for Lys9 methylation (H3K9me), which is found in transcriptionally silent heterochromatin. To determine the functions of K9 modification on variant versus canonical H3, we compared the phenotypes caused by engineering *H3.3*^*K9R*^ and *H3*^*K9R*^ mutant genotypes in *Drosophila melanogaster*. Whereas most *H3.3*^*K9R*^ and a small number of *H3*^*K9R*^ mutant animals are capable of completing development and do not have substantially altered protein coding transcriptomes, all *H3.3*^*K9R*^ *H3*^*K9R*^ combined mutants die soon after embryogenesis and display decreased expression of genes enriched for K9ac. These data suggest that the role of K9ac in gene activation during development can be provided by either H3 or H3.3. Conversely, we found that H3.3K9 is methylated at telomeric transposons, and this mark contributes to repressive chromatin architecture, supporting a role for H3.3 in heterochromatin that is distinct from that of H3. Thus, our genetic and molecular analyses demonstrate that K9 modification of variant and canonical H3 have overlapping roles in development and transcriptional regulation, though to differing extents in euchromatin and heterochromatin.

## INTRODUCTION

DNA interacts with histones and other proteins to establish chromatin environments that affect all DNA-dependent processes. The establishment of chromatin environments is accomplished through multiple mechanisms that collectively comprise the bulk of epigenetic regulation found in eukaryotes. In particular, post-translational modification (PTM) of histones influences DNA/histone interactions and also provides binding sites for recruitment of chromatin modulators that influence gene expression, DNA replication and repair, and chromosome segregation during cell division (Wallrath et al. 2014). In addition to histone PTMs, epigenetic regulation is modulated by the type of histone protein deposited onto DNA. There are two major categories of histone proteins: the canonical histones and the closely related histone variants (Talbert and Henikoff 2010, 2017). These two histone categories are distinguished by the timing of their expression during the cell cycle and their mechanism of deposition onto DNA. Canonical histones are encoded by multiple genes (e.g., ~55 in humans and ~500 in flies), organized into clusters that are highly expressed during S-phase of the cell cycle, and are deposited onto DNA by the histone chaperone CAF-1 in a replication-coupled manner (Marzluff et al. 2002; Tagami et al. 2004; Verreault et al. 1996). In contrast, variant histones are typically encoded by one or two genes, are expressed throughout the cell cycle, and can be deposited onto DNA independently of replication by histone chaperones other than CAF-1 (Henikoff and Ahmad 2005; Tagami et al. 2004; Szenker et al. 2011). Variant histones are often deposited at specific genomic locations and have functions that can differ from canonical histones. For example, two histone H2A variants, H2AX and H2A.Z, play critical roles in DNA repair (Scully and Xie 2013; Price and Andrea 2014), and the histone H3 variant CENP-A localizes to centromeres and is essential for kinetochore formation (Blower and Karpen 2001; Henikoff and Ahmad 2005; Mellone and Allshire 2003).

The major histone H3 variant in animal genomes is H3.3, which in both mice and *Drosophila* is encoded by two different genes (*H3.3A* and *H3.3B*) that produce identical proteins. Variant histone H3.3 differs from canonical H3.2 and H3.1 by only four or five amino acids, respectively (Szenker et al. 2011). In each case, three of these different amino acids are located in the globular domain of H3.3 and are necessary and sufficient for interaction with the replication-independent chaperones HIRA and ATRX-DAXX (Tagami et al. 2004; Goldberg et al. 2010; Ahmad and Henikoff 2002; Lewis et al. 2010). In H3.2, the only replication-dependent histone in *Drosophila*, the fourth amino acid difference occurs at position 31 in the unstructured N-terminal tail (Szenker et al. 2011). Histones H3.2 and H3.1 (collectively hereafter referred to as H3) along with H3.3 are some of the most conserved proteins in all eukaryotes (Malik and Henikoff 2003).

The conservation of amino acid differences between H3 and H3.3 during evolution strongly suggests that these proteins perform distinct functions. Indeed, H3.3 and H3 are deposited in different genomic regions in a variety of species (Mito et al. 2005; Schwartz and Ahmad 2005; Tamura et al. 2009; Jin et al. 2011; Kraushaar et al. 2013; Allis and Wiggins 1984). H3.3 is also enriched for different histone PTMs than H3 (Hake et al. 2006; McKittrick et al. 2004), and H3.3 containing nucleosomes can be less stable than those with H3 (Jin and Felsenfeld 2007; Xu et al. 2010). Although the epigenetic PTM signature on variant and canonical H3 histones is distinct, the degree to which particular histone PTMs found on both H3 and H3.3 can compensate for one another is not fully understood. Here, we explore the common and distinct functions of variant and canonical H3K9 function during *Drosophila* development.

H3.3 is associated with transcriptionally active regions of the genome with high nucleosome turnover, consistent with H3.3 being enriched in “activating” histone PTMs and depleted in “repressing” histone PTMs (Hake et al. 2006; McKittrick et al. 2004). One of the histone PTMs enriched on H3.3 relative to H3 is acetylation of lysine nine (K9ac), a mark associated with accessible chromatin (Hake et al. 2006; McKittrick et al. 2004). Previous studies have identified K9ac at promoters of genes and in regions of high transcriptional activity (Kharchenko et al. 2011; Bernstein et al. 2005; Liang et al. 2004; Roh et al. 2005). Additionally, mutation of H3K9 acetyltransferases results in compromised transcriptional output, suggesting K9ac contributes to or is a consequence of gene expression activation (Wang et al. 1998; Georgakopoulos and Thireos 1992; Kuo et al. 1998). Importantly, H3K9 acetyltransferases target other histone residues and have non-histone substrates as well (Glozak et al. 2005; Spange et al. 2009), indicating that one cannot deduce the function of K9ac solely by mutation of H3K9 acetyltransferases. For example, whereas mutation of the H3K9 acetyltransferase Rtt109 in budding yeast results in sensitivity to DNA-damaging agents, H3K9R mutants, which cannot be acetylated by Rtt109, are insensitive to DNA-damaging agents (Fillingham et al. 2008). Direct investigation of K9ac function in vivo therefore requires mutation of H3K9 itself. Previously, we used a *Drosophila* histone gene replacement platform (McKay et al. 2015) to generate a canonical *H3*^*K9R*^ mutant, and found no significant changes in gene expression at regions of the genome enriched in K9ac (Penke et al. 2016). This observation raises the possibility that H3.3K9ac functions in gene regulation and can compensate for the absence of H3K9ac.

H3.3 is also found at transcriptionally inactive, heterochromatic regions of the genome (Goldberg et al. 2010; Lewis et al. 2010; Wong et al. 2010). Heterochromatin is enriched in H3K9 di- and tri-methylation (me2/me3), modifications that recruit Heterochromatin Protein 1 (HP1) and are essential for heterochromatin function (Bannister et al. 2001; Lachner et al. 2001; Nakayama et al. 2001; Penke et al. 2016). DNA within heterochromatin is composed of repeated sequence elements, many of which are transcriptionally silent and consist of immobile transposons or transposon remnants. Using H3.3 mutants it was recently demonstrated that H3.3 is essential for repression of endogenous retroviral elements and that H3.3 can be methylated at lysine nine (Elsässer et al. 2015). H3.3K9me3 is also important for heterochromatin formation at mouse telomeres (Udugama et al. 2015). These studies did not assess the contribution of canonical H3K9 because strategies for mutating all replication-dependent H3 genes in mammalian cells have not been developed. We recently showed in *Drosophila* that mutation of canonical H3K9 causes defects in heterochromatin formation and transposon repression (Penke et al. 2016), similar to phenotypes observed in *C. elegans* in the absence of H3K9 methyltransferases (Zeller et al. 2016). In addition, we detected low levels of K9me2/me3 in *H3*^*K9R*^ mutants. Combined, these data suggest methylated H3.3K9 plays a role in heterochromatin formation and can compensate for the absence of canonical H3K9. However, the extent of functional overlap between variant and canonical H3K9 and the intriguing possibility that identical modifications on variant or canonical histones have distinct functions has yet to be fully investigated.

In order to better understand the functions of H3 and H3.3 and to compare the functions of the variant and canonical H3K9 residues, we used CRISPR-Cas9 to generate a variant K9R substitution mutation (*H3.3*^*K9R*^) in *Drosophila* and combined this with our previously described canonical *H3*^*K9R*^ mutant (Penke et al. 2016). By comparing the individual mutant phenotypes of *H3.3*^*K9R*^ and *H3*^*K9R*^ to the combined *H3.3*^*K9R*^ *H3*^*K9R*^ mutants using a variety of genomic and cell biological assays, we demonstrate that variant and canonical versions of H3K9 can compensate for each other, although to substantially different extents in euchromatin versus heterochromatin. H3K9 plays a more substantial role than H3.3K9 in heterochromatin formation and in the repression of transposons, whereas they compensate for each other in controlling euchromatic gene expression, particularly in regions enriched in the activating modification, K9ac.

## MATERIALS AND METHODS

### Generation of K9R mutant genotypes

Variant *H3.3*^*K9R*^ mutants generated by the cross scheme illustrated in Supplementary Figure 1A were selected by the absence of GFP fluorescence and/or the presence of straight wings. 1^st^ instar larvae from the variant and canonical *H3.3^K9R^ H3.2*^*K9R*^ cross described in Supplementary Figure 1B were selected based on the presence of GFP fluorescence. Only larvae that receive the H3^HWT^ or H3^K9R^ transgene will survive embryogenesis, as this transgene provides the only source of canonical histone genes. In Table 2, rows one and two indicate progeny from the cross *yw; H3.3^2x1^ / CyO, twiGFP x yw; Df(2L)BSC110 / CyO, twiGFP*. Rows three and four indicate progeny from the cross *H3.3B^K9R^; H3.3^2x1^ / CyO, twiGFP x H3.3B^K9R^; Df(2L)BSC110 / CyO, twiGFP*. In these crosses, the expected ratio of heterozygous to homozygous *H3.3A*^*2x1*^ animals is 2:1, as *CyO, twiGFP/CyO, twiGFP* animals do not eclose as adults.

**Figure 1:**
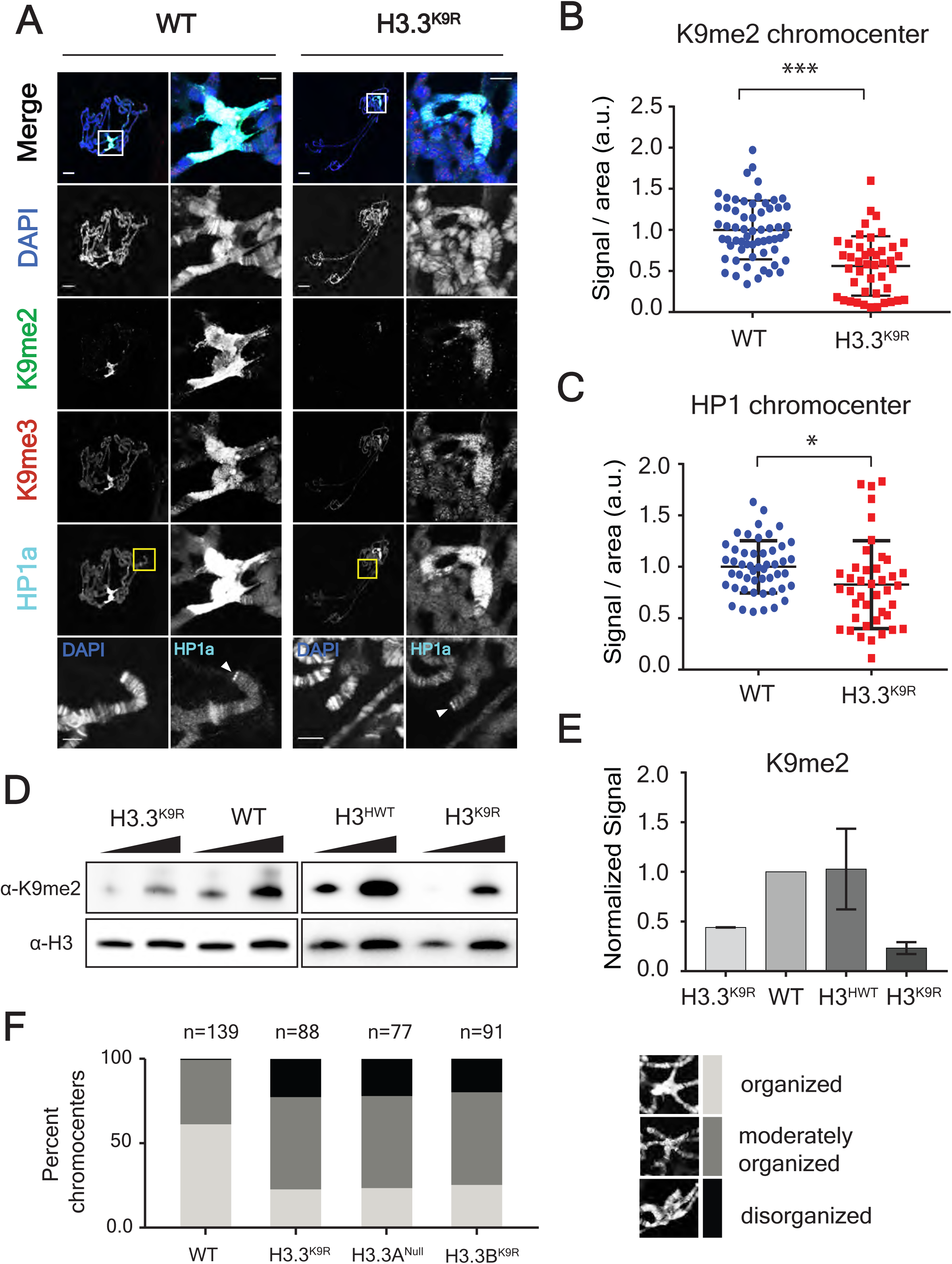
K9me2/me3 and HP1a signal is decreased in H3.3^K9R^ mutants. A) 3^rd^ instar larval salivary gland polytene chromosome spreads from wild-type (left) and H3.3^K9R^ mutants (right) stained with anti-K9me2, anti-K9me3, anti-HP1a, and DAPI to mark DNA. Right panel for each genotype shows enlarged chromocenter indicated by white boxes. Bottom panel shows magnified view of telomere indicated by yellow boxes. Scale bar = 20 microns (whole polytene) 5 microns (chromocenter/telomere). B, C) Immunofluorescent signal of K9me2 (B) or HP1a (C) at chromocenters in wild-type (WT) and H3.3^K9R^ mutants (a.u. = arbitrary units). Values were normalized to area of the chromocenter and set relative to the average WT value from matched slides (see Supplementary Experimental Methods). Significance was determined using t-test (* p<0.05, ** p<0.005, *** p<0.0005). D) Western blot of K9me2 from salivary glands with H3 used as loading control. E) K9me2 signal was quantified by densitometry and normalized to corresponding H3 loading control band. Normalized values were set relative to WT normalized signal. Error bars represent standard error of the mean from two independent biological replicates (see Materials and Methods). F) Quantification of chromocenter organization from *WT, H3.3^K9R^, H3.3A^Null^, H3.3B*^*K9R*^ mutants.

### CRISPR-Cas9 Mutagenesis and Transgene Integration

A single gRNA targeting *H3.3B* near the K9 residue was inserted into pCFD3 and co-injected with a 2 kb homologous repair template containing the H3.3BK9R substitution (Supplementary Experimental Methods). Constructs were injected into embryos expressing Cas9 from the *nanos* promoter (*nanos-cas9*; Kondo and Ueda 2013). Recovered *H3.3B*^*K9R*^ alleles were subsequently crossed into *H3.3A* null backgrounds (*H3.3A*^*2x1*^ over deficiency *Df(2L)BSC110*). Independent *H3.3B*^*K9R*^ CRISPR alleles were used to generate trans-heterozygous animals for all experiments. To generate *H3.3B* rescue constructs, a 5 kb genomic sequence containing the entire wild-type *H3.3B* transcription unit was PCR amplified from genomic DNA of *nanos-cas9* flies and cloned into pATTB (Supplementary Experimental Methods). Gibson assembly (Gibson et al. 2009) using primers containing K9R or K9Q substitutions was used to generate mutated versions of *H3.3B*, and all three constructs were integrated into the 86FB *attP* landing site by ΦC31-mediated recombination.

### Immunofluorescence

Salivary gland preparations stained using anti-H3K9me2, anti-H3K9me3, anti-H3K9ac, or anti-HP1a were performed as previously described (Cai et al. 2010). Antibody sources and concentrations are included in Supplementary Experimental Methods. 1^st^ instar larval brains were prepared similar to imaginal wing disc preparations described in Estella et al. (2008).

### Western Blots

ImageJ densitometry analysis was used to determine K9me2, K9ac, or H3 band intensity (See Supplementary Experimental Methods). Histone modification signal was normalized to corresponding H3 loading control signal. Normalized signal from different titrations of the same genotype were averaged and consequent values were set relative to WT value. This process was completed for two biological replicates for both K9me2 and K9ac.

### Sample Preparation and Sequence Data Analysis

FAIRE-seq and RNA-seq samples were prepared from wandering 3^rd^ instar imaginal wing discs as previously described (McKay and Lieb 2013). Sequencing reads were aligned to the dm6 (6.04) reference genome using Bowtie2 (FAIRE) and Tophat (RNA) default parameters (Langmead and Salzberg 2012; Trapnell et al. 2014). FAIRE peaks were called with MACS2 using a shift size of 110bp and a stringency cutoff of 0.01 (Zhang et al. 2008). Transcripts were assembled with Cufflinks (Trapnell et al. 2014). Bedtools was used to determine read coverage at peaks and transcripts (Quinlan and Hall 2010) and DESeq2 was used to determine statistical significance (p<0.05) (Love et al. 2014). The following modENCODE 3^rd^ instar larval ChIP-seq data sets were used: K9me2=GSE47260, and K9me3=GSE47258. K9ac ChIP-seq data from imaginal wings discs was generated by Pérez-Lluch et al. (GSM1363590, 2015).

Chromatin state analysis was performed using data from Kharchenko et al (2010), which assigns small regions of the genome into one of nine different chromatin state. FAIRE peaks were classified as one or more chromatin states based on overlap with regions defined by Kharchenko et al. (2010). Of all the peaks in a particular chromatin state, we determined the percentage of peaks that had significantly different FAIRE signal in mutant compared to WT samples. RNA chromatin state analysis was performed in a similar fashion.

See Supplemental Experimental Procedures for a detailed description of the methods. Strains are available upon request. Sequencing data are available at GEO under accession number GSE106192.

## RESULTS

### H3.3^K9R^ mutant animals are viable but sterile

In order to investigate the role of H3.3K9 in *Drosophila* development and compare it to the role of H3K9, we first generated an H3.3^K9R^ animal by introducing a K9R substitution at the endogenous *H3.3B* locus using CRISPR/Cas9 and then combining recovered *H3.3B*^*K9R*^ mutant alleles with a previously generated *H3.3A* null allele (H3.3A/B combined genotype denoted hereafter as *H3.3*^*K9R*^; see Table 1, Table S1, and Figure S1A for histone genotype nomenclature) (Sakai et al. 2009). These *H3.3*^*K9R*^ mutants, which contain the full complement of endogenous canonical *H3* genes, eclose as adults at the expected Mendelian ratios (Table 2) and appear morphologically normal. Therefore, canonical H3 can provide all of the H3K9 function during *Drosophila* development. This result is consistent with a previous study finding that flies without any H3.3 protein could be propagated as a stock if canonical H3.2 was expressed from a transgene using the *H3.3B* promoter (Hödl and Basler 2012). Our results are also in line with a previous report in which *H3.3A* and *H3.3B* null animals containing an *H3.3A*^*K9R*^ transgene were viable (Sakai et al. 2009). However, whereas these *H3.3A*^*K9R*^ transgenic animals were fertile (Sakai et al. 2009), we found that animals with an endogenous *H3.3B*^*K9R*^ mutation and the same *H3.3A* null allele used by Sakai et al. (2009) were sterile. The sterility of our *H3.3*^*K9R*^ animals was rescued in both males and females by a transgene containing the wild-type *H3.3B* gene ectopically integrated into the genome, suggesting that the relative abundance of H3.3^K9R^ causes sterility (Figure S2). We conclude that H3.3K9 plays an essential role during gametogenesis and speculate that different amounts of H3.3^K9R^ histones from *H3.3A* or *H3.3B* promoters may account for the differences between our observations and those of Sakai et al. (2009).

**Table 1:** Genotype description of H3.3 and H3 K9R mutants.

| Nomenclature<sup>a</sup> | Canonical Histone Genotypes |  | Variant Histone Genotypes |  |
| --- | --- | --- | --- | --- |
|  | Endogenous | Transgenic | H3.3B | H3.3A |
| *WT* | WT | - | WT | WT |
| *H3.3B<sup>K9R</sup>* | WT | - | K9R | WT |
| *H3.3A<sup>Null</sup>* | WT | - | WT | Δ |
| *H3.3<sup>K9R</sup>* | WT | - | K9R | Δ |
| *H3<sup>HWT</sup>* | Δ | WT | WT | WT |
| *H3<sup>K9R</sup>* | Δ | K9R | WT | WT |
| *H3.3<sup>K9R</sup> H3<sup>HWT</sup>* | Δ | WT | K9R | Δ |
| *H3.3<sup>K9R</sup> H3<sup>K9R</sup>* | Δ | K9R | K9R | Δ |

<sup>a</sup> Wild-type (WT), gene deletion (Δ), no transgenic histone array (-). See Supplementary Table 1 for full genotypes.

**Table 2:** *H3.3*^*K9R*^ mutants are viable but sterile.

| *H3.3B* | *H3.3A<sup>a</sup>* | Observed | Expected | p<sup>b</sup> | Fertile |
| --- | --- | --- | --- | --- | --- |
| *WT* | Δ/+ | 535 | 535.3 | n.s. | yes |
| *WT* | Δ | 268 | 267.7 | n.s. | yes |
| *K9R* | Δ/+ | 400 | 438 | <0.005 | yes |
| *K9R* | Δ | 257<sup>c</sup> | 219 | <0.005 | No<sup>d</sup> |

<sup>a</sup> *H3.3A<sup>2x1</sup>* deletion allele
<sup>b</sup> p-value calculated with a chi-square test
<sup>c</sup> The higher than expected number of observed *H3.3B<sup>K9R</sup> H3.3A<sup>Null</sup>* animals is presumably due to non-specific detrimental effects caused by the presence of a balancer chromosome in siblings with the *H3.3B<sup>K9R</sup>* mutation and balancer-derived wild-type *H3.3A*.
<sup>d</sup> Both males and females.

### H3.3K9 and H3K9 have overlapping functions during development

We previously observed that canonical *H3*^*K9R*^ mutants could complete development, although 98% of these mutant animals died during larval or pupal stages (Penke et al. 2016). We considered the possibility that *H3*^*K9R*^ mutant animals progressed to late larval or pupal stages of development because of compensation by H3.3K9. We therefore tested if the *H3.3*^*K9R*^ genotype would advance the *H3*^*K9R*^ mutant stage of lethality by observing the development of animals in which the *H3.3*^*K9R*^ and *H3*^*K9R*^ mutant genotypes were combined (Table 1, Table S1, and Figure S1B). The *H3*^*K9R*^ genotype was generated using our previously described histone replacement platform (McKay et al. 2015; Penke et al. 2016). Briefly, the endogenous array of ~100 canonical histone gene clusters was deleted and replaced with an ectopically located transgene encoding a BAC-based, tandem array of 12 canonical histone gene clusters in which the *H3* genes contain a K9R mutation (Figure S1B). A 12x tandem array of wild-type, canonical histone genes (denoted <u>h</u>istone <u>w</u>ild type or *H3*^*HWT*^, Figure S1B), which fully rescues deletion of the endogenous histone gene array, was used as a control (McKay et al. 2015). Similar to the *H3.3*^*K9R*^ mutants, *H3*^*HWT*^ animals with the *H3.3*^*K9R*^ mutant genotype (denoted hereafter as *H3.3^K9R^ H3^HWT^*; see Table 1 and Figure S1B) were viable (Table 3). However, only 34.6% of *H3.3*^*K9R*^ *H3*^*HWT*^ progeny eclosed as adults (Table 3) compared to essentially 100% of the *H3.3*^*K9R*^ genotype that contained the full complement of endogenous, wild-type *H3* genes (Table 2). This result suggests that in the presence of fewer total canonical *H3* gene copies, the *H3.3*^*K9R*^ mutation is more detrimental. Importantly, animals with the *H3.3^K9R^ H3*^*K9R*^ combined mutant genotype containing both the variant and canonical K9R mutation were 100% inviable, dying with high penetrance at the 1^st^ instar larval stage, much earlier than the majority of *H3*^*K9R*^ mutants. These results demonstrate that H3.3K9 can partially compensate for the absence of H3K9, indicating that H3.3K9 and H3K9 have some redundant functions.

**Table 3:**
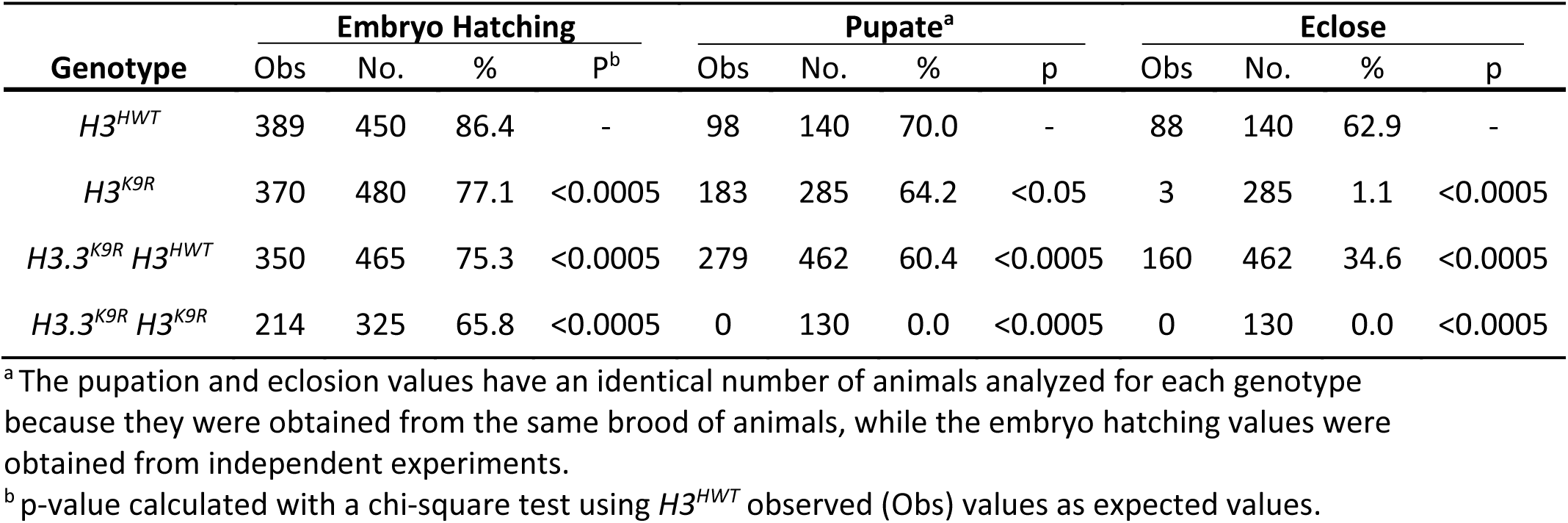
*H3.3*^*K9R*^ and *H3*^*K9R*^ mutations are synthetically lethal.

### H3K9 PTMs are lost in animals lacking H3.3K9 and H3K9

We previously found that K9me2/me3 signal in *H3*^*K9R*^ mutant animals is substantially reduced but not absent. Thus, a possible reason why *H3.3^K9R^ H3*^*K9R*^ mutants have a more severe developmental defect than *H3*^*K9R*^ mutants is complete loss of K9me throughout the genome. We therefore assessed K9me2/me3 levels in *H3.3*^*K9R*^ and *H3.3^K9R^ H3*^*K9R*^ mutants by immunofluorescence. We first assessed K9me2/me3 levels in salivary gland polytene chromosomes of *H3.3*^*K9R*^ mutants, with the expectation that if H3.3K9 is methylated the signal will be reduced relative to controls. The salivary gland is a highly polyploid tissue (>1000C) and the alignment of chromatids in the polytene chromosomes results in easily visible structures that provide information about levels and genomic locations of histone PTMs using immunofluorescence. *H3.3*^*K9R*^ mutants had lower levels of both K9me2 and K9me3 compared to wild-type controls at the largely heterochromatic chromocenter, demonstrating that H3.3K9 is normally methylated in the pericentric heterochromatin of otherwise wild-type animals (Figure 1A, B). In support of this result, western blot analysis of salivary glands demonstrated that K9me2 levels were decreased in *H3.3*^*K9R*^ mutants compared to wild-type controls (Figure 1D, E).

Because *H3.3*^*K9R*^ mutants exhibited reduced K9me2/me3 signal at the chromocenter, we next used immunofluorescence to examine localization of HP1a, which binds K9me2/me3. In line with reduced K9me2/me3 signal, HP1a signal at the chromocenter of H3.3^K9R^ mutants was reduced compared to wild-type controls (Figure 1A, C). HP1a and H3.3 also both localize to telomeres (Goldberg et al. 2010; Lewis et al. 2010). We found that HP1a localizes to telomeres in *H3.3*^*K9R*^ mutants (Figure 1A), as it does in *H3*^*K9R*^ mutants (Penke et al. 2016). These results are consistent with previous observations that HP1 recruitment to telomeres requires telomere binding proteins (Raffa et al. 2011; Vedelek et al. 2015; Badugu et al. 2003) and not the H3K9 methyltransferase Su(var)3-9 (Perrini et al. 2004), suggesting that H3K9me is not required for HP1 recruitment to telomeres.

Because *H3.3^K9R^ H3*^*K9R*^ combined mutants do not develop to the 3^rd^ instar larval stage, we examined K9me2 levels in 1^st^ instar larval brains. *H3*^*K9R*^ mutants (with wild-type variant histones) and *H3.3*^*K9R*^ *H3*^*HWT*^ mutants (with a 12x transgenic complement of wild-type canonical histone genes) each exhibited reduced K9me2 levels by immunofluorescence compared to *H3*^*HWT*^ controls, consistent with the polytene chromosome data (Figure 2A). In contrast, the *H3.3^K9R^ H3*^*K9R*^ variant and canonical combined mutant brains had undetectable levels of K9me2 in the vast majority of cells (Figure 2A). These results provide further evidence that H3.3K9 is methylated and that the total amount of K9me is derived from both H3.3 and H3.

**Figure 2:**
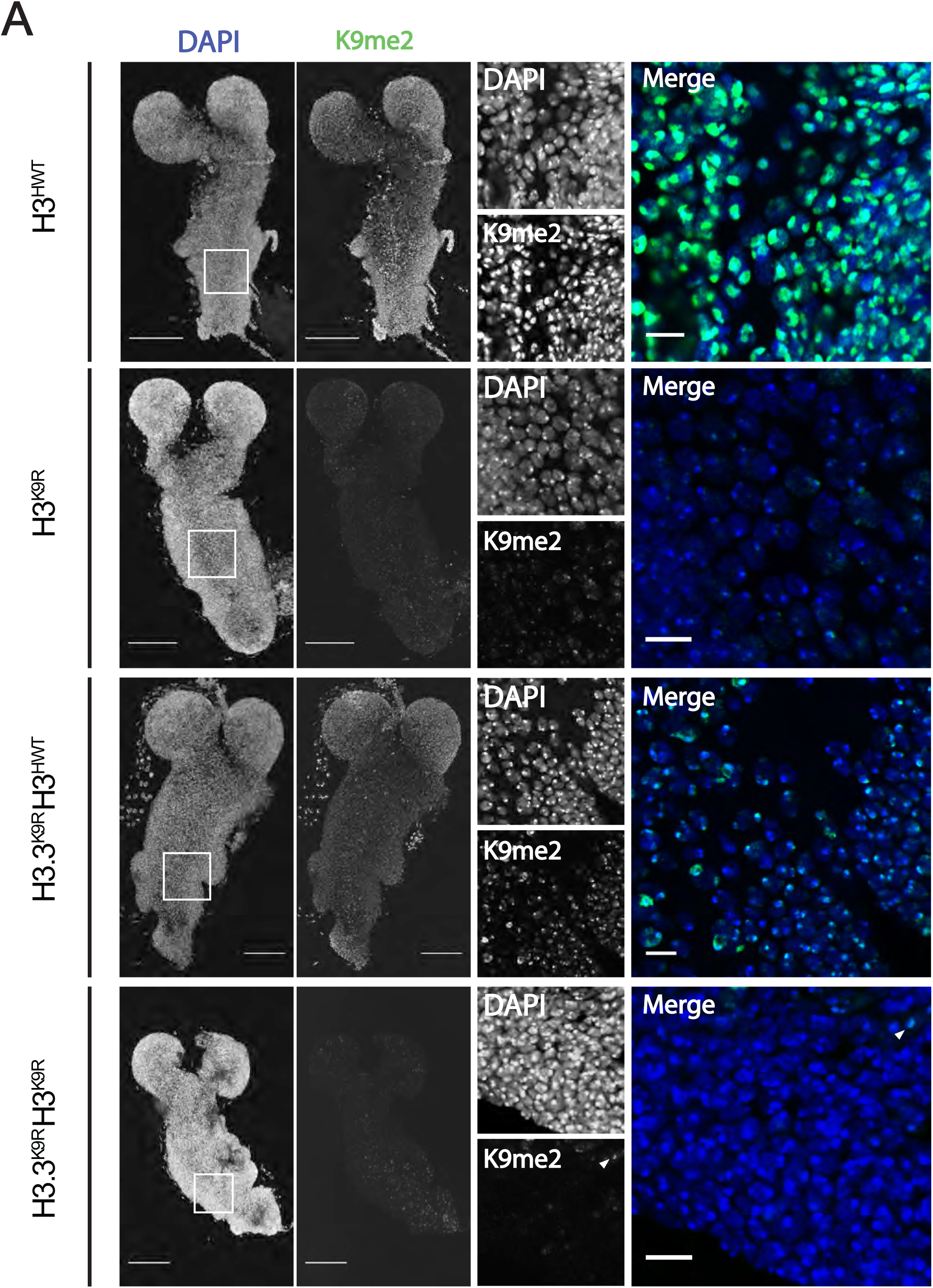
K9me2/me3 signal is diminished in K9R mutants. A) 1^st^ instar larval brains stained with anti-K9me2 and DAPI to mark DNA from *H3*^*HWT*^, *H3*^*K9R*^, *H3.3^K9R^ H3^HWT^*, and *H3.3^K9R^ H3*^*K9R*^ animals. Left panel shows max projection of 2 micrometer confocal sections through the entire brain. Right panel shows a magnified, single confocal section from the area indicated by the white boxes. Arrowheads indicate cells with residual K9me2 signal in *H3.3^K9R^ H3*^*K9R*^ animals. Scale bar = 50 microns (whole brain) 10 microns (enlarged image).

Interestingly, a small number of cells in the *H3.3^K9R^ H3*^*K9R*^ 1^st^ instar mutant brains retained low levels of K9me2 signal at the chromocenter (arrowheads, Figure 2). Cells with residual K9me2 express ELAV, a pan-neuronal marker, and lack expression of Deadpan and Prospero, markers of proliferating neuroblasts and ganglion mother cells, respectively (circles, Figure S3). These data indicate that cells with K9me2 positive chromocenters in *H3.3^K9R^ H3*^*K9R*^ mutant 1^st^ instar larval brains are differentiated neurons. We suspect that the K9me2 signal in these cells reflects maternally provided wild-type H3 protein remaining in the genomes of quiescent neurons that differentiated prior to having their maternal H3 fully replaced by zygotically expressed H3K9R mutant histones. A corollary to this conclusion is that the proliferating neuroblasts and their GMC daughters likely have progressed through a sufficient number of S phases such that replacement of maternal H3 with zygotic H3K9R eliminates detectable K9me2 signal.

We also found that levels of H3K9 acetylation were reduced in both the *H3.3*^*K9R*^mutant and the *H3*^*K9R*^ mutant relative to controls, as determined both by immunofluorescence of salivary gland polytene chromosomes (Figure 3A, B) and by western blots of salivary gland extracts (Figure 3C). Because a substantial amount of K9ac is placed on H3.3, we considered the possibility that lack of K9ac was responsible for the fertility defects of *H3.3*^*K9R*^ mutants and the early lethality of *H3.3^K9R^ H3*^*K9R*^mutants. To address this question, we integrated either an H3.3B^K9^, an H3.3B^K9R^, or an H3.3B^K9Q^ transgene into the same genomic position in order to determine if a K9Q acetyl mimic could restore fertility to *H3.3*^*K9R*^ mutants. Animals with only an *H3.3B*^*K9R*^ mutation at the endogenous locus (i.e., containing a wild-type *H3.3A* gene), and carrying either an H3.3B^K9R^ or H3.3B^K9Q^ transgene were sterile, precluding us from constructing the genotype to test if these transgenes could rescue the sterility of *H3.3*^*K9R*^ mutant adults (Figure S2). This result suggests that both the H3.3B^K9R^ and H3.3B^K9Q^ transgenes acted dominantly to compromise fertility. Furthermore, these data imply that H3.3B^K9R^ and H3.3B^K9Q^ histones are incorporated into chromatin.

**Figure 3:**
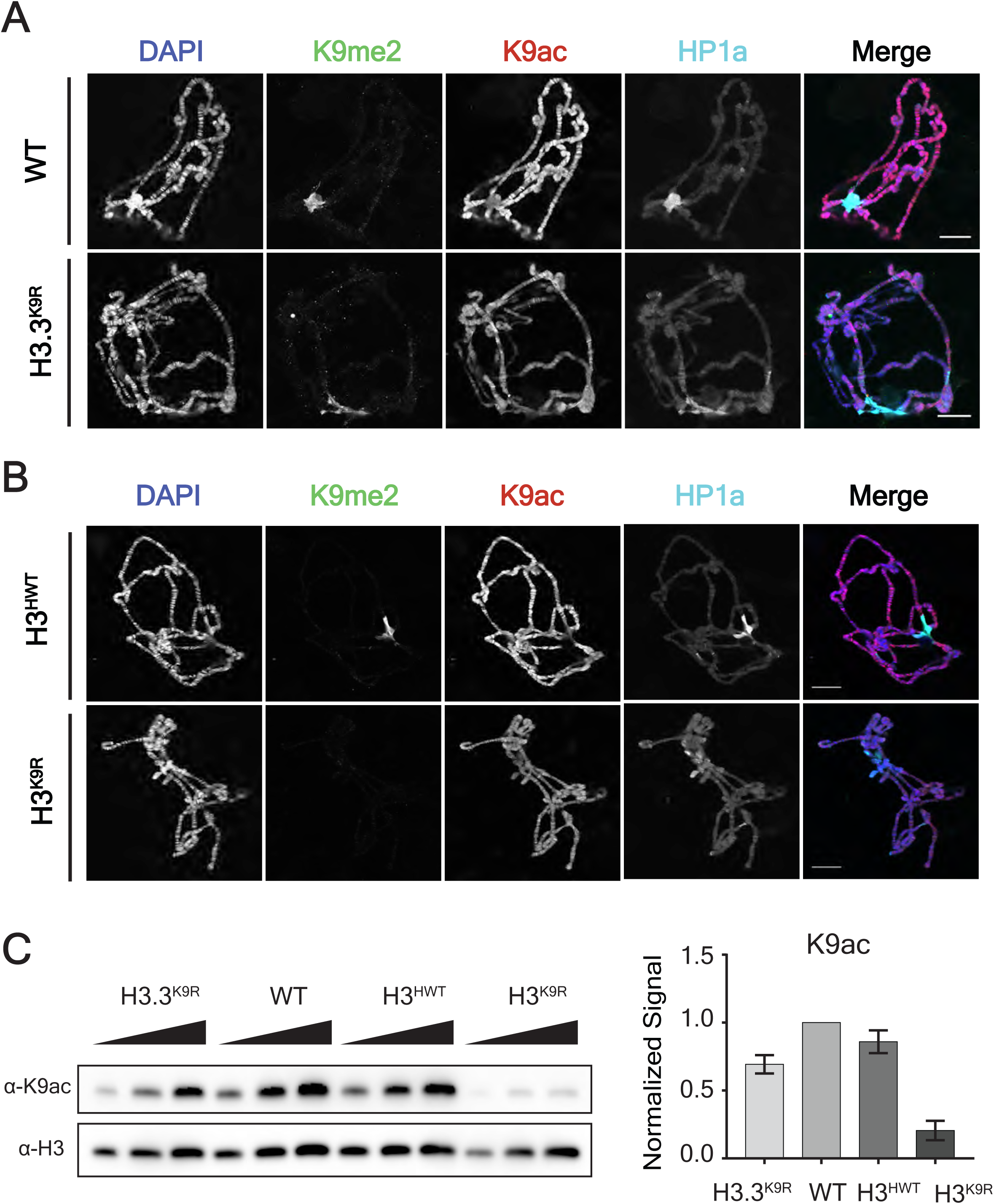
K9ac signal is decreased in H3.3^K9R^ mutants. A,B) Polytene chromosome spreads from wild-type (*WT*) and *H3.3*^*K9R*^ mutants (A) or *H3*^*HWT*^ and *H3*^*K9R*^ mutants (B) stained with anti-K9me2, anti-K9ac, anti-HP1a, and DAPI to mark DNA. Scale bar = 20 microns. C) Western blot of K9ac from salivary glands with H3 used as loading control. K9ac signal was quantified by densitometry and normalized to corresponding H3 loading control band. Normalized values were set relative to WT normalized signal. Error bars represent standard error of the mean from two independent biological replicates (see Materials and Methods).

### H3.3K9 regulates chromatin organization at the chromocenter, telomeres, and transposons

We next asked if the reduction of K9me2/me3 in *H3.3*^*K9R*^ mutants affected chromatin organization by cytological examination of salivary gland polytene chromosomes using DAPI staining of DNA. As we found previously in *H3*^*K9R*^ mutants (Penke et al. 2016), in some *H3.3*^*K9R*^ mutants polytene chromosome spreads the chromocenter appeared abnormal and not fully condensed (Figure 1F). The cause of this phenotype is unclear but may reflect altered chromatin organization or defects in the under-replication of salivary gland pericentric heterochromatin (Belyaeva et al. 1998; Zhimulev et al. 2003). Based on their cytology, we binned chromocenters into three categories: “organized”, “moderately organized”, and “disorganized” (Figure 1F). We categorized chromocenters from four genotypes: wild-type (WT; i.e., with the endogenous canonical histone genes), an *H3.3A* null mutant (*H3.3A^Null^*), an *H3.3B* K9R substitution mutant (*H3.3B*^*K9R*^), and the *H3.3B*^*K9R*^; *H3.3A*^*Null*^ double mutant in which all H3.3 contains the K9R substitution (*H3.3*^*K9R*^) (Table 1, Table S1, and Figure S1A). Whereas the majority of wild-type chromocenters were organized (60% organized vs 40% moderately organized), both the *H3.3B*^*K9R*^ and the *H3.3A*^*Null*^ single mutants had increased percentages of moderately organized and disorganized chromocenters (Figure 1F). For example, ~22% of chromocenters in the various *H3.3* mutants were disorganized compared to less than 1% of wild-type chromocenters. These results indicate that H3.3 contributes to chromocenter structure. Interestingly, the *H3.3B*^*K9R*^; *H3.3A*^*Null*^ double mutant had the same proportion of moderately organized and disorganized chromocenters as either single mutant. This result suggests that either reducing *H3.3* gene dose (i.e., the *H3.3A*^*Null*^ allele) or expressing K9R mutant H3.3 histones (i.e., the *H3.3B*^*K9R*^ mutation), can prevent normal H3.3 function at pericentric heterochromatin.

Given the disrupted chromocenter structure in *H3.3*^*K9R*^ mutants, we next examined chromatin structure genome wide by performing Formaldehyde Assisted Isolation of Regulatory Elements followed by whole genome sequencing (FAIRE-seq). FAIRE-seq provides a measure of local nucleosome occupancy across the genome, revealing regions of “open” chromatin that are relatively depleted of nucleosomes (Simon et al. 2013). We previously found using this technique that regions of heterochromatin enriched in K9me, particularly pericentromeric heterochromatin, were more open in canonical *H3*^*K9R*^ mutants relative to *H3*^*HWT*^ controls (Penke et al. 2016). To determine if variant *H3.3*^*K9R*^ mutants had a similar phenotype we performed FAIRE-seq in triplicate on imaginal wing discs from wandering 3^rd^ instar larvae in *WT*, *H3.3A^Null^*, *H3.3B*^*K9R*^, and *H3.3B*^*K9R*^; *H3.3A^Null^ (H3.3^K9R^)* double mutant genotypes. Sequencing reads were aligned to the genome and peaks were called on each of the three replicates and combined into a merged peak set. Called peaks were consistent across replicates and read coverage across peaks was highly correlated (R≥ 0.96) (Figure S4A, B). Additionally, wild-type FAIRE data was consistent with previously generated data from wing discs (McKay and Lieb 2013) (Figure S4D). *H3.3A^Null^, H3.3B^K9R^, and H3.3*^*K9R*^ mutants each had a similar percentage of peaks with significantly altered FAIRE signal when compared to wild-type: 8.8%, 6.5%, and 7.9% respectively (Figures 4A-C). Moreover, significantly changed peaks across the three mutants exhibited a high degree of overlap. Of the 2,660 significantly changed peaks across all mutants, 21% were shared among all three and 52% by at least two mutants (Figure S5A). FAIRE signal at significantly changed peaks also displayed similar fold changes in mutants compared to wild-type and were not exacerbated in the double mutant compared to either single mutant (Figure S5B). These data suggest H3.3A and H3.3BK9 both function to regulate chromatin architecture.

**Figure 4:**
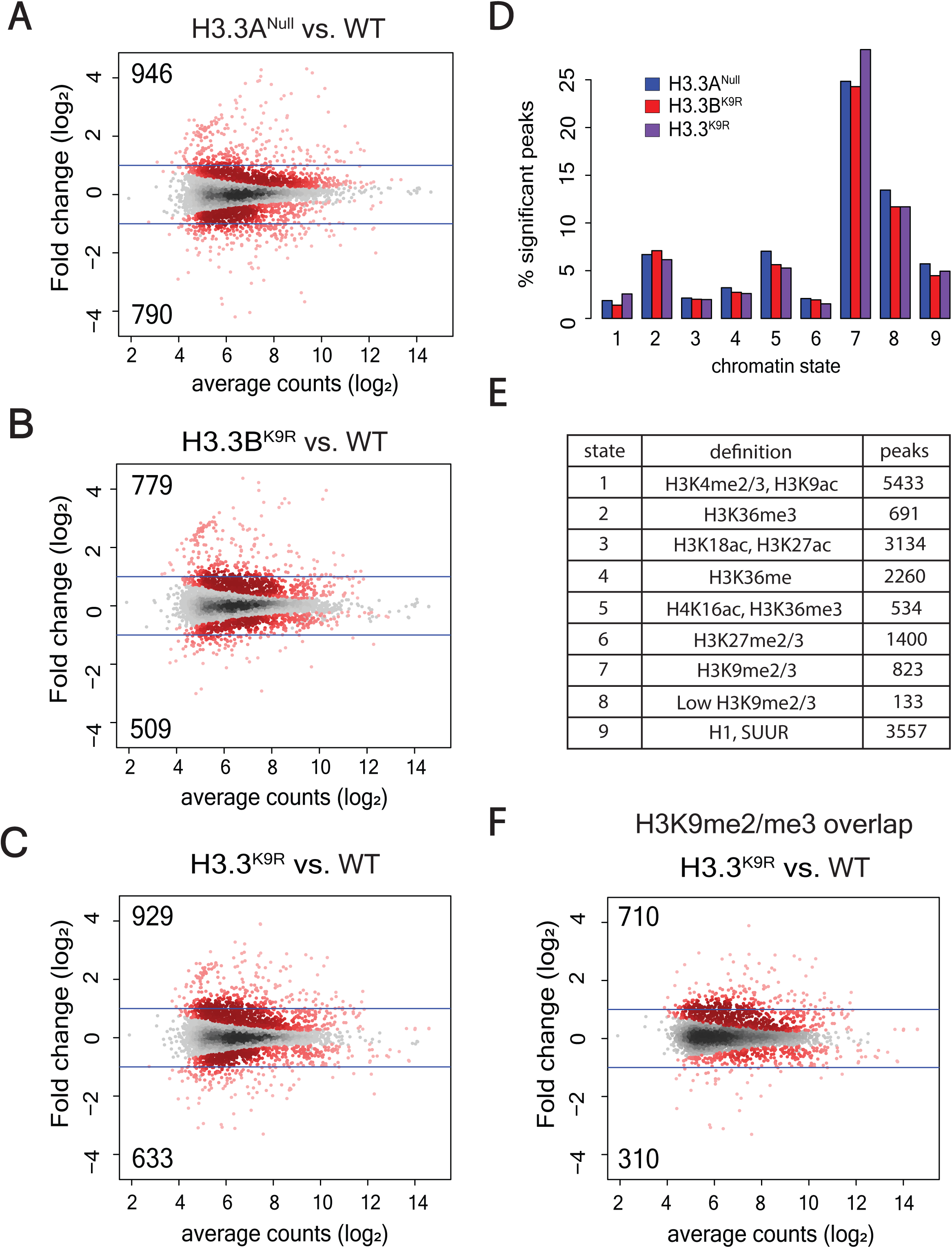
H3.3K9 regulates chromatin architecture in regions of K9me. A-C) Mutant: WT ratio of *H3.3A*^*null*^ (A), *H3.3B*^*K9R*^ (B), or *H3.3*^*K9R*^ (C) FAIRE signal from 3^rd^ instar imaginal wing discs at 19,738 FAIRE peaks called by MACS2. Red dots indicate significantly different peaks (p<0.05), and insets indicate the number of significantly increased (top) or decreased (bottom) peaks. Average counts signify average normalized reads that overlap a peak in mutant and WT samples. D) Percentage of peaks in a particular chromatin state that have significantly different FAIRE signal in mutants versus WT (top). Bottom panel shows a summary of histone modifications or proteins that define a chromatin state and the number of FAIRE peaks assigned to a given chromatin state. E) Boxplot of FAIRE enrichment over input at 126 transposon families (* indicates p < 0.05 and *** indicates p <0.0005). F) Plot from C showing only those peaks that overlap an K9me2 or K9me3 peak from modENCODE ChIP-seq data.

We next asked if the changes in FAIRE signal we observed in *H3.3* mutants were characterized by a particular chromatin signature. We assigned each called FAIRE peak to one of nine different chromatin states characterized by different combinations of histone PTMs as defined by Kharchenko et al. (2010). We then calculated the percentage of FAIRE peaks that changed between an *H3.3* mutant and wild-type within each chromatin state. Regions of K9me2/me3 showed the highest percentage of changes in FAIRE signal in *H3.3A^Null^*, *H3.3B*^*K9R*^, and the *H3.3*^*K9R*^ mutant compared to wild-type, supporting the idea that H3.3K9 is methylated and plays a necessary role in regulating chromatin architecture (Figure 4D). Changes in FAIRE signal were also more likely to occur in regions of H3K36me3, a mark that is enriched along gene bodies that are themselves enriched for H3.3 (Bannister et al. 2005; Szenker et al. 2011). Finally, we used modENCODE K9me2 and K9me3 ChIP-seq data to complement the chromatin state analysis. Of the FAIRE peaks significantly increased or decreased in *H3.3*^*K9R*^ mutants compared to wild-type, 76.4% and 49.0% respectively overlapped a K9me2 or K9me3 peak (Figure 4F). These results demonstrate that altered FAIRE signal in *H3.3*^*K9R*^ mutants occurred in regions normally occupied by K9me.

We also observed increased FAIRE signal at telomeres in all three *H3.3* mutant genotypes, particularly on chromosomes 2R and 3L (Figure S5C), suggesting that H3.3 regulates telomeric chromatin architecture. In *Drosophila,* telomeres are composed of retrotransposons enriched in K9me2/me3 (Levis et al. 1993; Cenci et al. 2005). H3.3 plays a similar role in the mouse, in which H3.3 null mutant embryonic stem cells exhibit an increase in transcripts from transposons (Elsässer et al. 2015) and telomeres (Udugama et al. 2015). Additionally, we previously observed transposon activation and mobilization in canonical *H3*^*K9R*^ mutants (Penke et al. 2016). For these reasons, we examined FAIRE signal at transposons in our *H3.3* mutants using the piPipes pipeline, which avoids ambiguity in aligning reads to repetitive transposons by mapping to transposon families (Han et al. 2015). Both *H3.3A*^*Null*^ and *H3.3B*^*K9R*^ mutants resulted in significantly increased FAIRE signal at transposons, and *H3.3*^*K9R*^ mutants had on average even higher increased FAIRE signal at transposons (Figure 5A, B). Moreover, FAIRE signal at some telomeric transposons, particularly TART-B, was increased in *H3.3* mutants (Figure 5C). However, the extent of increase in *H3.3*^*K9R*^ mutants was not as severe as previously observed for *H3*^*K9R*^ mutants (Penke et al. 2016) (Figure 5B). These results support a role for H3.3K9 in chromatin-mediated transposon repression, though to a lesser extent than H3K9.

**Figure 5:**
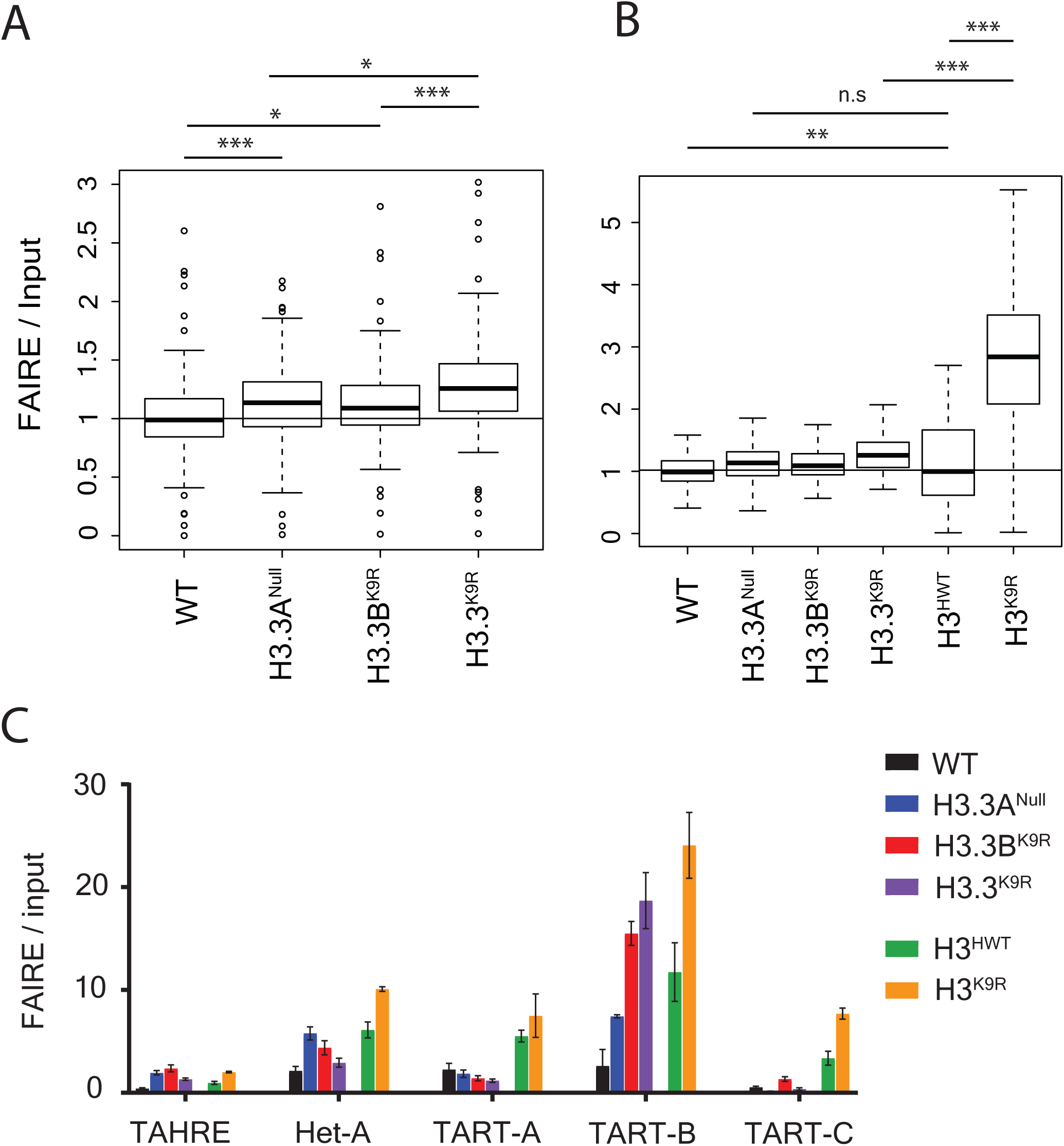
Imaginal wing disc FAIRE signal of *H3.3* mutants is increased at telomeres and transposons. A) Boxplot of average FAIRE enrichment determined by piPipes pipeline across 126 transposon families (Han et al. 2015). Genomic DNA from *Drosophila* embryos used as input control. B) Boxplots in A shown alongside FAIRE enrichment for *H3*^*HWT*^ and *H3*^*K9R*^ mutants from a separate experiment (Penke et al. 2016). C) FAIRE enrichment of *H3.3* and *H3*^*K9R*^ mutants at telomeric transposons. Error bars indicate standard deviation from three replicates for each genotype. Statistical significance determined by paired t-test (p<0.05 *, p<0.005 **, p<0.0005 ***, n.s. = not significant).

### H3.3K9 and H3K9 functions overlap in regions of K9ac and partially in regions of K9me

To investigate the cause of lethality when both variant and canonical H3 histones contain the K9R mutation, we performed RNA-seq of 1^st^ instar larvae from four genotypes: *H3*^*HWT*^, *H3*^*K9R*^, *H3.3^K9R^ H3^HWT^*, and *H3.3^K9R^ H3*^*K9R*^ (Table 1, Table S1). Larvae of the correct genotype were identified by GFP fluorescence (see Materials and Methods). RNA sequencing reads were aligned to the genome using Tophat, transcript assembly was performed by Cufflinks, and DESeq2 was used for statistical analysis (Trapnell et al. 2014; Love et al. 2015). Each genotype was verified by examination of RNA-seq reads mapping to the K9 codon of variant and canonical histones. Correlation analysis demonstrated transcript abundance across all assembled transcripts was highly similar among replicates, and was also similar to previously generated data from wild-type 1^st^ instar larvae (Figure S6) (Graveley et al. 2011). Additionally, histone expression was similar across all genotypes, suggesting that variation in histone levels do not underlie observed phenotypes (Figure S7A). In line with our previous analysis of *H3*^*K9R*^ RNA-seq data from imaginal wing discs (Penke et al. 2016), the majority of significantly changed transcripts in *H3*^*K9R*^ 1^st^ instar samples was increased compared to *H3*^*HWT*^ (247 increased vs 41 decreased), supporting a role for H3K9me in gene silencing (Figure 6A). *H3.3*^*K9R*^ *H3*^*HWT*^ samples had a similar number of significantly changed transcripts, and again most transcripts showed increased signal compared to *H3*^*HWT*^ (203 vs 126), though fold changes were smaller than *H3*^*K9R*^ mutants (Figure 6B). By contrast, the *H3.3^K9R^ H3*^*K9R*^ combined mutant genotype caused a much more pronounced effect on gene expression compared to either the *H3.3*^*K9R*^ *H3*^*HWT*^ or the *H3*^*K9R*^ mutant genotypes (Figure 6C); 869 transcripts exhibited increased RNA signal and 1036 transcripts were decreased compared to *H3*^*HWT*^ samples. The number of decreased transcripts in *H3.3^K9R^ H3*^*K9R*^ animals compared to *H3*^*HWT*^ was therefore about ten-fold higher than either the variant or canonical K9R mutant alone. Thus, similar to our viability analysis (Table 3), these RNA-seq results demonstrated that variant and canonical versions of H3K9 compensate for each other in the regulation of gene expression.

**Figure 6:**
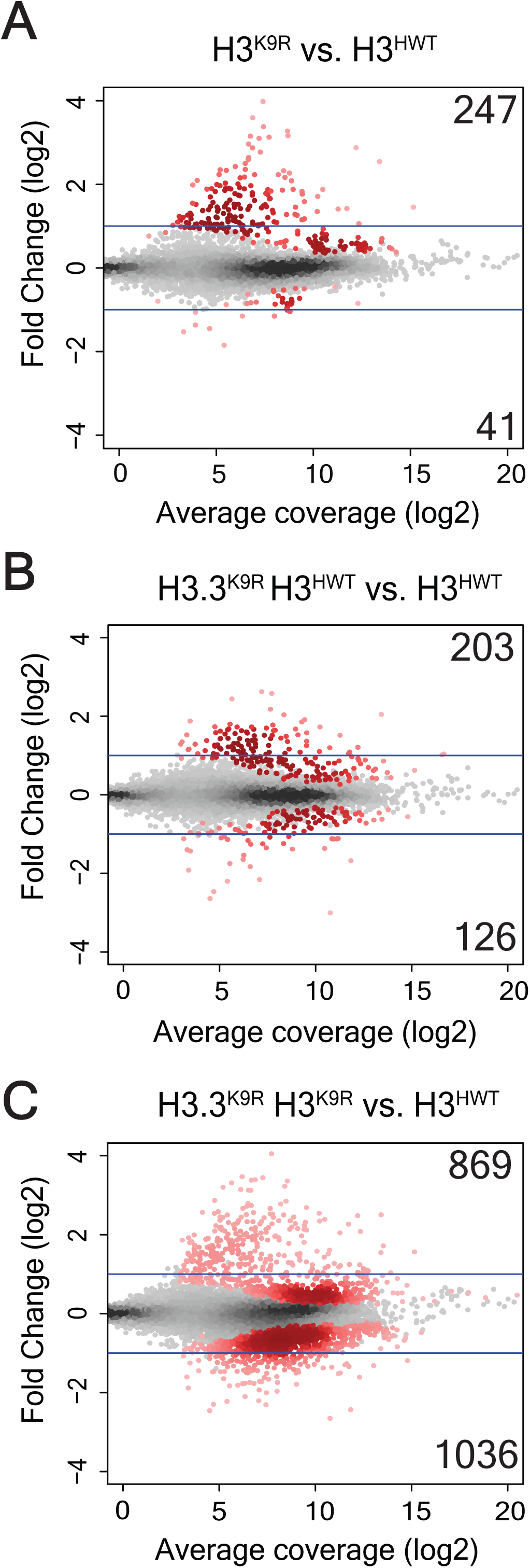
H3.3K9 and H3K9 redundantly regulate gene expression. A-C) Mutant: WT ratio of *H3*^*K9R*^ (A), *H3.3*^*K9R*^ (B), or *H3.3^K9R^ H3*^*K9R*^ (C) RNA signal from 1^st^ instar larvae at 10,253 transcripts assembled by Cufflinks. The Y axis indicates the log_2_ transformation of mutant/control signal between the genotypes being compared (indicated at the top of each plot). Red dots indicate significantly different transcripts (p<0.05) and insets signify the number of significantly increased (top) or decreased (bottom) transcripts. Average coverage signifies the average number of normalized reads that overlap a transcript in mutant and *H3*^*HWT*^ samples.

Because we observed increases in FAIRE signal at transposons in *H3.3*^*K9R*^ mutants from wing disc samples, we examined RNA levels of transposon families in 1^st^ instar larvae. Similar to our previous RNA-seq observations from *H3*^*K9R*^ mutant wing discs (Penke et al. 2016), RNA signal at transposons in *H3*^*K9R*^ 1^st^ instar larvae was increased relative to the *H3*^*HWT*^ control (Figure S7B, C). Although on average transposon levels were only slightly higher in *H3.3*^*K9R*^ *H3*^*HWT*^ mutants compared to *H3*^*HWT*^, transposon levels in *H3.3^K9R^ H3*^*K9R*^ combined mutants were significantly higher than either *H3.3*^*K9R*^ *H3*^*HWT*^ or *H3*^*K9R*^ mutants alone (Figure S7B, C). Moreover, telomeric transposons are generally increased in all K9R mutants compared to *H3*^*HWT*^ controls (Figure S7D). Together these results support an overlapping role for H3.3K9 and H3K9 in regulating gene expression and transposon repression.

We next examined chromatin signatures of significantly altered transcripts to explore the mechanism of the observed gene expression changes. All transcripts were assigned to one or more chromatin states based on their overlap with genomic regions defined by Kharchenko et al. (2010). We then determined the percentage of transcripts within a given chromatin state that were either increased or decreased in K9R mutants relative to *H3*^*HWT*^ controls (Figure 7 A-C). Transcripts in regions of K9me2/me3 (chromatin state 7 and 8) were the most likely to have significantly increased RNA levels in mutants compared to *H3*^*HWT*^. Although *H3.3^K9R^ H3*^*K9R*^ combined mutants had the highest percentage of chromatin state 7 transcripts that were significantly increased (~26%), *H3*^*K9R*^ mutants also displayed a high percentage (~13%) of change within chromatin state 7 (Figure 7A, D, Figure S8A). These results suggest that H3.3K9 contributes to gene repression in regions of K9me2/me3 but cannot completely compensate for the absence of H3K9.

**Figure 7:**
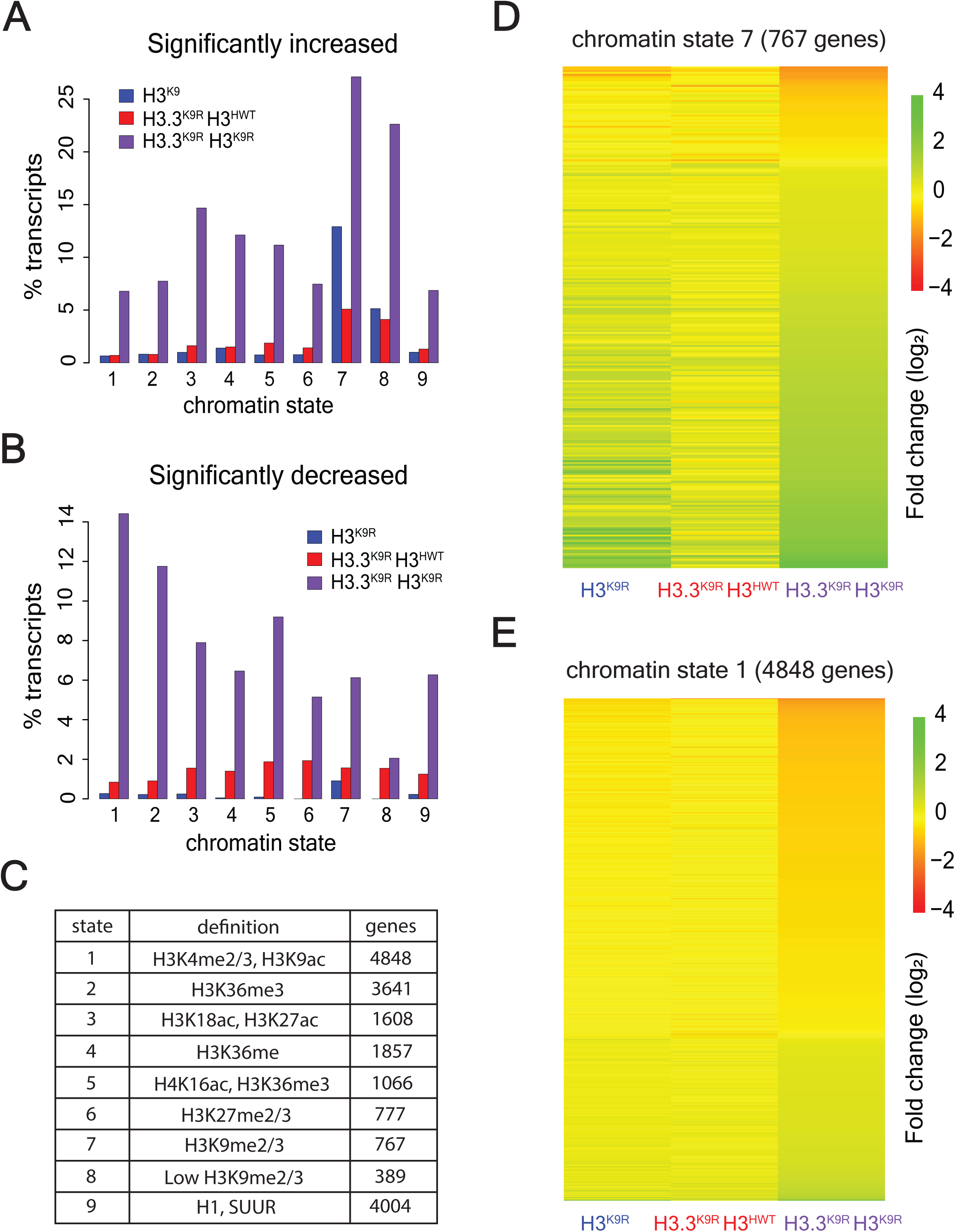
H3.3K9 and H3K9 redundancy differs in heterochromatin and euchromatin. A, B) Percentage of transcripts in a chromatin state that have significantly increased (A) or decreased (B) RNA signal in mutants versus *H3*^*HWT*^. C) Table indicates the number of transcripts that overlap a particular chromatin state. D, E) Heatmaps showing fold change of K9R mutants over *H3*^*HWT*^ at chromatin state 7 regions (D) and chromatin state 1 regions (E). Each row indicates a transcript that overlaps the indicated chromatin state.

In contrast to upregulated transcripts, very few transcripts were significantly decreased in *H3.3*^*K9R*^ *H3*^*HWT*^ or *H3*^*K9R*^ mutants. However, the *H3.3^K9R^ H3*^*K9R*^ combined mutant displayed numerous significant decreases in transcript abundance. Interestingly, transcripts in chromatin state 1, characterized by K9ac and lack of K9me, were most likely to be decreased (Figure 7B, E). Several other chromatin states showed elevated transcript changes, particularly in the *H3.3^K9R^ H3*^*K9R*^ combined mutant; however, in this analysis transcripts can be assigned to more than one chromatin state. Indeed, many transcripts in chromatin state 1 also overlap other chromatin states. We therefore performed a supplementary analysis that examined only transcripts that overlap a single chromatin state. This analysis demonstrated that transcripts solely in chromatin state 1 were much more likely to change in K9R mutants than those in other chromatin states (Figure S8A). Similar results were obtained using imaginal wing disc K9ac ChIP data from Pérez-Lluch et al. (2015). Whereas few transcripts that overlapped K9ac were significantly altered in either single mutant (68 in *H3*^*K9R*^ and 116 in *H3.3^K9R^ H3^HWT^*), 1195 K9ac associated transcripts exhibited changed expression levels in *H3.3^K9R^ H3*^*K9R*^ combined mutants (Figure 8). These data suggest that in regions of K9ac, H3.3 and H3 can completely compensate for each other. Additionally, these data provide evidence that K9ac facilitates gene expression.

**Figure 8:**
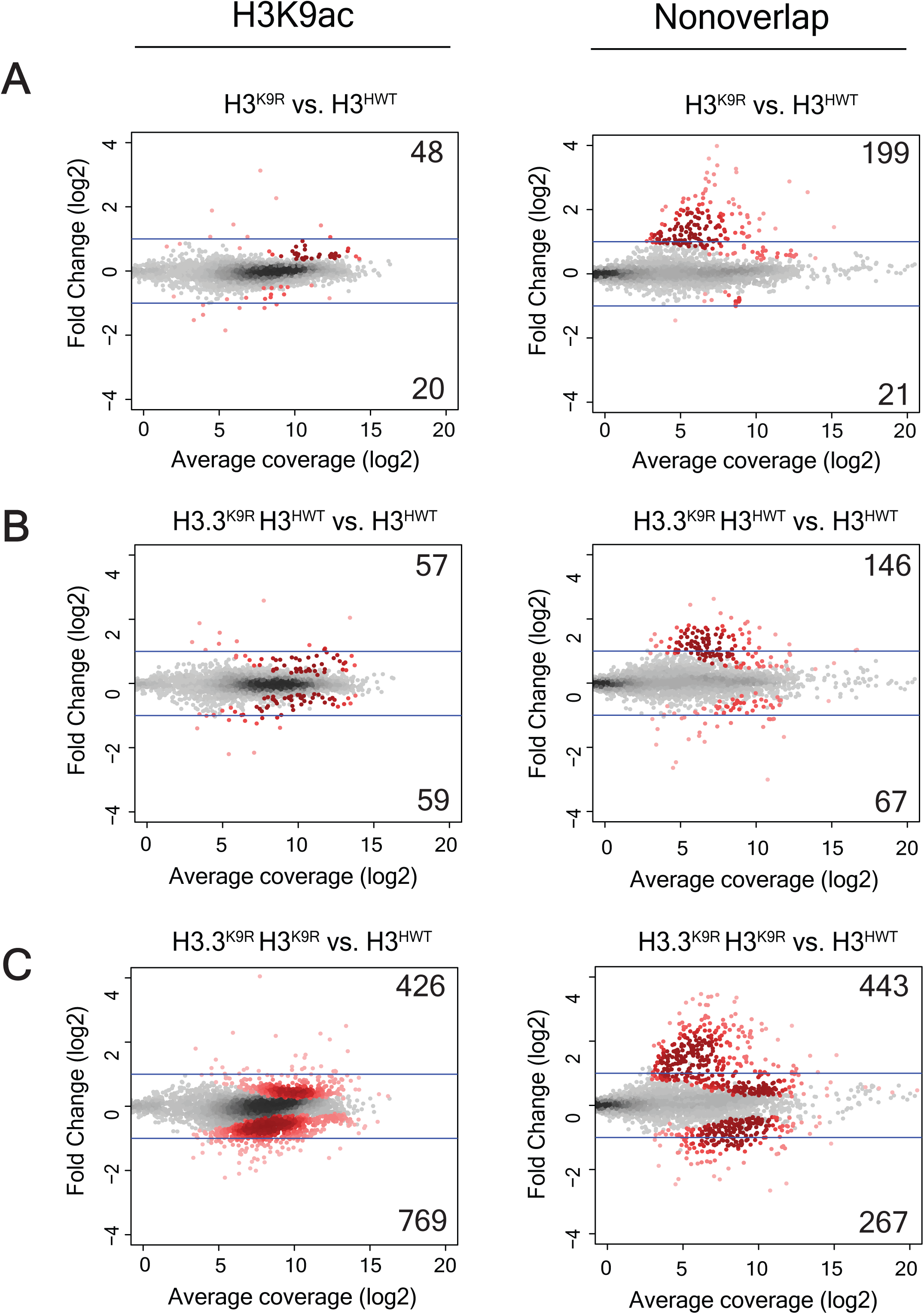
K9ac associated transcripts are altered in *H3.3^K9R^ H3*^*K9R*^ double mutants. MA plot showing fold change of normalized RNA signal in *H3*^*K9R*^ (A), *H3.3*^*K9R*^ *H3*^*HWT*^ (B), and *H3.3^K9R^ H3*^*K9R*^ (C) mutants versus *H3*^*HWT*^ at all transcripts from merged transcriptome. Average coverage on X-axis represents the mean expression level of a transcript. Transcripts that overlap an K9ac peak called from ChIP-seq data (GSM1363590; Pérez-Lluch et al. 2015) are shown in the left panel while those that do not are shown in the right panel. Significance (shown in red) was determined using DESeq2 (Love et al. 2015) and an adjusted p value cutoff of 0.05.

## DISCUSSION

### Overlapping and distinct developmental functions of H3 and H3.3

In this study, we determined the distinct and overlapping roles that lysine 9 of variant and canonical histone H3 play in gene expression and heterochromatin function during *Drosophila* development. Our developmental genetic analyses demonstrate that H3.3K9 is necessary for fertility but not viability in *Drosophila*. In addition, we find that some euchromatic functions of H3K9 can be provided by either variant H3.3 or canonical H3, whereas H3.3K9 cannot completely compensate for H3K9 in some regions of heterochromatin as discussed below.

Several studies from multiple species have investigated the developmental functions of H3.3 and H3. In mice, single mutation of either *H3.3A* or *H3.3B* results in reduced viability and compromised fertility (Bush et al. 2013; Couldrey et al. 1999). Similarly, *Drosophila H3.3A* and *H3.3B* double mutants appear at lower than expected Mendelian ratios and are sterile (Sakai et al. 2009). H3.3 in *Tetrahymena thermophila* is also important for sexual reproduction, although it is not required for viability or maintenance of nucleosome density (Cui et al. 2006). Both *Tetrahymena* and *Drosophila* H3.3 and H3 can compensate for one another. In *Tetrahymena*, canonical H3 is dispensable if H3.3 is overexpressed (Cui et al. 2006). Similarly in *Drosophila*, transgenic expression of H3 can rescue both the semi-lethality (Sakai et al. 2009) and infertility (Hödl and Basler 2012) of *H3.3* mutants, indicating some functional redundancy between the two histones. Indeed, when expressed equivalently, *Drosophila* H3.3 can provide all of the developmental functions of H3 (Hödl and Basler 2012). Moreover, H3.3 is the sole H3 protein in *S. pombe* and *S. cerevisiae* yeast (Malik and Henikoff 2003).

### H3.3K9 functions in heterochromatin

We find that under endogenous expression conditions, H3.3K9 functions in heterochromatin, including pericentromeric and telomeric regions of the genome. We detected H3.3K9 methylation in pericentromeric heterochromatin, congruous with previous data demonstrating that H3.3 in *Drosophila* is deposited at the chromocenter of polytene chromosomes in a replication-dependent manner (Schwartz and Ahmad 2005). We also observed that *H3.3*^*K9R*^ mutants exhibited an abnormal chromocenter structure in polytene chromosomes. Moreover, we provide evidence that H3.3K9 is required for maintenance of telomeric chromatin architecture and repression of certain telomeric transcripts, indicating that replication-coupled expression of H3 cannot provide these particular H3K9 functions. These findings in *Drosophila* are consistent with studies in mouse embryonic stem cells showing that H3.3 is localized to telomeres, is methylated at K9, and functions in repression of telomeric repeat-containing RNAs (Goldberg et al. 2010; Udugama et al. 2015). Conversely, the genetic data we presented here and previously (Penke et al. 2016) indicate that H3K9 is essential for repression of transposon-derived transcripts in pericentric heterochromatin, and H3.3K9 cannot compensate for the lack of H3K9 at these regions of the genome. The role of H3.3K9 in telomere structure and function may be independent of HP1, as HP1 is recruited to telomeres via the terminin complex independently of H3K9me (Raffa et al. 2011; Vedelek et al. 2015; Badugu et al. 2003).

### K9ac regulates euchromatic gene expression

Previous studies that mapped histone modifications across the genome identified K9ac as a characteristic of transcriptionally active regions (Kharchenko et al. 2011; Bernstein et al. 2005; Liang et al. 2004; Roh et al. 2005). Moreover, mutation of H3K9 acetyltransferases results in compromised transcriptional activity (Wang et al. 1998; Georgakopoulos and Thireos 1992; Kuo et al. 1998). However, H3K9 acetyltransferases have non-histone substrates in addition to H3K9, and decreased transcriptional output may be the result of pleiotropic effects (Glozak et al. 2005; Spange et al. 2009; Fillingham et al. 2008). Our study provides evidence that K9ac, rather than non-histone targets of H3K9 acetyltransferases, contributes to activating transcription, as *H3.3*^*K9R*^ and *H3*^*K9R*^ mutants exhibit reduced gene expression in regions normally enriched for K9ac. Importantly, these K9ac rich regions with reduced gene expression are not normally enriched in K9me2 or me3, indicating the observed phenotype is not due to changes in K9me2 or me3 and likely results from loss of K9ac. This change in gene expression was accompanied by a fully penetrant lethality early in larval development of *H3.3^K9R^ H3*^*K9R*^ combined mutant animals, raising the possibility that gene expression control via acetylation of H3K9 is critical for the completion of animal development. These data are also consistent with previous studies in *C. elegans* demonstrating that H3K9 methylation is not essential for viability (Towbin et al. 2012; Zeller et al. 2016).

### Overlapping and distinct genomic functions of H3K9 and H3.3K9

Functional overlap of H3K9 and H3.3K9 appears to vary at different regions of the genome. Whereas H3.3K9 and H3K9 can perform similar functions in euchromatic regions of the genome and can fully compensate for one another, our RNA-seq data demonstrate H3.3K9 can only partially compensate for H3K9 in regions of heterochromatin. Partial compensation by H3.3K9 in regions of K9me2/me3 is in line with previous studies showing H3.3 is found at heterochromatin (Goldberg et al. 2010; Lewis et al. 2010; Wong et al. 2010) and plays a role in transposon repression (Elsässer et al. 2015). In the genotypes we analyzed, mRNA encoding variant and canonical H3 are expressed from their native promoters. Thus, disparity in functional overlap might be due to differences in modes of expression and deposition and thus total amounts of variant and canonical H3 histones in particular regions of the genome. For instance, H3 is normally enriched in heterochromatin compared to H3.3 (Ahmad and Henikoff 2002), which may cause *H3*^*K9R*^ mutations to be more detrimental in these regions. However, H3.3 may be able to provide all H3 function when highly expressed in a replication-dependent manner, as a transgenic histone gene array in which the *H3.2* coding region was replaced by *H3.3* is nearly fully functional in larval imaginal discs (Hödl and Basler 2012). Thus, differences in expression and/or deposition into chromatin may be the only basis for functional differences between H3.3 and H3.2 that we observed.

Heterochromatin may be particularly sensitive to incorporation of non-modifiable K9 residues. H3K9 methylation serves as a binding site for the protein HP1, which can in turn recruit H3K9 methyltransferases (Elgin and Reuter 2013; Grewal and Jia 2007). Methylation of a neighboring nucleosome can restart the cycle and initiate propagation of a heterochromatic configuration along the chromosome. Introduction of even a small number of H3K9R containing nucleosomes may therefore disrupt this cycle and prevent proper heterochromatin formation and gene repression. Incorporation of H3.3B^K9R^ histones into regions of heterochromatin may dominantly affect chromatin structure, resulting in the observed phenotypes at pericentromeres and telomeres in H3.3^K9R^ mutants. In contrast, incorporation of low amounts of H3K9R histones in euchromatin may not reduce K9ac levels sufficiently to disrupt gene expression. Finally, amino acid differences in variant and canonical H3 may direct distinct histone modification states on different histone types by influencing the binding of chromatin modifying enzymes (Jacob et al. 2014). Different histone modification states on H3.3 and H3 may underlie variation in compensation at different genomic regions.

In sum, our data investigating H3.3K9 and H3K9 function provide evidence that K9ac activates gene expression and advance our understanding of the overlapping and distinct functional roles of variant and canonical histones.

## Acknowledgement

We thank Bhawana Bariar for assistance generating the gRNA construct, Jeff Sekelsky for the pBlueSurf construct, and Kami Ahmad for providing the *H3.3A*^*2×1*^ flies. This work was supported by 5T32GM007092-39 and F31GM115194 to T.J.R.P and R01DA036897 to D.J.M, B.D.S, A.G.M, and R.J.D.

